# Holobiont Urbanism: sampling urban beehives reveals cities’ metagenomes

**DOI:** 10.1101/2020.05.07.075093

**Authors:** Elizabeth Hénaff, Devora Najjar, Miguel Perez, Regina Flores, Christopher Woebken, Christopher E. Mason, Kevin Slavin

**Affiliations:** NYU Tandon School of Engineering, Brooklyn, NY; Center for Urban Science and Progress, NYU, New York; MIT Media Lab, Cambridge, Mass; Parson’s School of Design, New York, NY; Extrapolation Factory, New York, NY; Department of Physiology and Biophysics, Weill Cornell Medicine, New York, NY, USA; The HRH Prince Alwaleed Bin Talal Bin Abdulaziz Alsaud Institute for Computational Biomedicine, Weill Cornell Medicine, New York, NY, USA; The WorldQuant Initiative for Quantitative Prediction, Weill Cornell Medicine, New York, NY, USA

## Abstract

Over half of the world’s population lives in urban areas with, according to the United Nations (UN), nearly 70% expected to live in cities by 2050 (United Nations, 2019). Our cities are built by and for humans, but are also complex, adaptive biological systems involving a diversity of other living species. The majority of these species are invisible and constitute the city’s microbiome. Our design decisions for the built environment shape these invisible populations, and we interact with them on a constant basis. A growing body of evidence shows us that our health and well-being are dependent on these interactions. Indeed, multicellular organisms owe meaningful aspects of their development and phenotype to interactions with the microorganisms—bacteria or fungi—with which they live in continual exchange and symbiosis. While the processing and sequencing of samples can be high-throughput, gathering samples is still very expensive, labor intensive, and can require mobilizing large numbers of volunteers to get a snapshot of the microbial landscape of a city, such as City Sampling Day (metasub.org). Here we postulate that honeybees may be effective collaborators in the sampling process, as they daily forage within a 2-mile radius of their hive. We describe the results of a pilot study conducted with 3 rooftop beehives in Brooklyn, NY, where we evaluated the potential of various hive materials (beeswax, honey, debris, pollen, propolis) to reveal information as to the surrounding metagenomic landscape, and where we conclude that the bee debris are the richest substrate. Based on these results, we profiled 4 additional cities in this manner: Sydney, Melbourne, Venice and Tokyo. While the molecular and computational methods used here were based on DNA analysis, it is possible they could be used to monitor RNA-based viruses such as Sars-Cov-2. Here we present the results of this study, and discuss them in terms of architectural implications, as well as the potential of this method for epidemic surveillance.

## 1. INTRODUCTION

Over half of the world’s human population lives in urban areas and, according to the U.N., nearly 70% of us will live in cities by 2050 (United Nations, 2019). Our cities are built by and for humans, but are also complex, adaptive biological systems involving a diversity of living species (Puppim de Oliveira et al. 2010). The majority of these species are invisible and constitute the city’s microbiome. Our design decisions for the built environment shape these invisible populations, and we interact with them on a constant basis. A growing body of evidence shows us that our health and well-being are dependent on these interactions. Indeed, multicellular organisms owe meaningful aspects of their development and phenotype to interactions with the microorganisms—bacteria or fungi—with which they live in symbiosis. Accumulated evidence confirms that mammalian phenotypes are a combination of an individual’s genetic makeup as well as its biome (microbiome and other organisms), including disease states such as obesity (Turnbaugh et al., 2006) and influence on neuro-psychiatric disorders as well (Cryan & Dinan, 2012). Beyond human consequences, plants’ flowering time has been found to depend on the soil microbiome (Wagner et al., 2015) and the useful metabolic compounds in medicinal plants are possibly synthesized in conjunction with their symbiont bacteria (Koberl, Schmidt, Ramadan, Bauer, & Berg, 2013), both traits were formerly thought to depend only on the plant’s genotype. Metagenomic studies such as these are facilitated by the rapidly decreasing cost of high-throughput DNA sequencing and this growing understanding that the phenotype of a multicellular organism depends on both its own genotype and that of its associated microbes leads us to redefine the scope of genetic identity. The holobiont, term which encompasses the notion of host and symbiont, and coined by Lynn Margulis in 1991 (Margulis and Fester 1991), can now be augmented to define the hologenome.

Metagenomics is a rapidly growing field that is well-situated to survey across all domains and kingdoms of life, including city-scale efforts of urban metagenomics. Microbial classification using high-throughput DNA sequencing is faster and more comprehensive than culture-based methods, and has enabled city-wide mapping of microbial populations (Afshinnekoo 2015, Hsu 2016, MetaSUB Consortium, 2016). Mapping the indoor environments (Lax 2017, Kembel 2014) also provides insights into the relationship between humans and the indoor microbiome, which holds promise for designing buildings that optimize this metric. Thus, we are moving away from the germ-centric paradigm of microbes to the quantification of a ubiquitous, continuous and commensal map of the environmental microbiome within which we live, work, and sleep. While the processing and sequencing of samples can be high-throughput (with automation, hundreds at a time), gathering samples is still very expensive, labor intensive, and can require mobilizing large numbers of volunteers to get a snapshot of the microbial landscape of a city, such as global City Sampling Day (metasub.org). Moreover, samples collected manually with swabs represent a limited area: 0.5-1m^2^. While this scale of resolution is important for applications such as tracking contamination through a hospital, it is not always easily implemented for city-scale studies and leads researchers to look for pinchpoints where samples might be most meaningful. Examples of this have been MetaSub sampling subways, air sampling in indoor environments (J. F. Meadow et al., 2014), or sewers (Maritz 2017).

Setting out to collect a more distributed and comprehensive sample of the urban landscape, following conversations with artists Timo Arnall and Jack Schulze, we investigated the potential of using honey bees as proxy sampling mechanisms for the urban microbiome. On average, honeybees forage within a 1-2 mile radius around their hive in rural environments (Eckhert 1933) and 0.3-1 miles in urban environments (Garbuzov, Schürch, and Ratnieks 2015), and we hypothesized that their travel would permit them to interact with various microbial environments including air, water, and mammalian sources in addition to their known plant targets. We designed a pilot study to test for geo-specific microbial residues corresponding to all of these environmental within material found in a hive.

Here we describe the results of a pilot study conducted with 3 rooftop beehives in Brooklyn, NY, where we evaluated the potential of various hive materials (beeswax, honey, debris, pollen, propolis) to reveal information as to the surrounding metagenomic landscape, and where we conclude that the bee debris are the richest substrate. Based on these results, we profiled 4 additional cities in this manner: Sydney, Melbourne, Venice and Tokyo. Here we present the results of this study, and discuss them in terms of architectural implications, as well as the potential of this method for epidemic surveillance.

## 2. RESULTS

### 2.1 Brooklyn Pilot study

In order to assess the potential of using honeybees as metagenomic “sample collectors”, we designed a pilot study with three Langstroth hives in Brooklyn, wherein we sampled the interior of the hive, the debris at the bottom, 3 bees from each hive, and its honey. We sequenced the DNA of each sample using a high-throughput shotgun approach, and classified the reads using DIAMOND-MEGAN against the NCBI NR nucleotide database, which includes all kingdoms and domains of life (see Methods for more details) (**Fig 1**). The honey of each hive is largely dominated by the species *Lactobacillus kunkeei* (**Fig 1A**), an obligate fructophilic lactic acid bacteria described as found in flowers, wine, and honey (Endo et al., 2012). Also of note are *Acinetobacter nectaris*, found in flowers (Álvarez-Pérez, Lievens, Jacquemyn, & Herrera, 2013), and *Zygosaccharomyces rouxii*, known to thrive under salt or sugar osmotic stress and thus cause food spoilage (Flagfeldt, Siewers, Huang, & Nielsen, 2009). Bee gut commensals were found in low abundance in honey, and include the species identified in the bee samples, described below. Traces of plant DNA were also identified, including *Medicago truncatula* and *Vitis vinifera*. The Bee samples (**Fig 1B**) contain sequences representative of both *Apis mellifera* (European honeybee) and *Apis dorsata* (Giant honeybee), indicating the hives are likely hybrids of these two species. The most abundant microbes in the bee samples include species described as bee commensals such as *Snodgrassella alvi* and *Gilliamella apicola* (Kwong & Moran, 2013), as well as *Lactobacillus wkB8* and *wkB10* (Kwong, Mancenido, & Moran, 2014). The bees from AS and FG hives display almost identical species distribution, however the CH sample shows lower abundances of the aforementioned commensals, and includes species absent from the other two. These include *Nosema cenarae*, a fungal parasite of the honeybee affecting both larvae and adults (Eiri, Suwannapong, Endler, & Nieh, 2015), as well as various human-related bacteria such as *Sporosarcina newyorkensis*, isolated from clinical samples in New York State (Wolfgang et al., 2012) and *Enterobacter* species. We hypothesize the colonization of atypical bacteria in this bee is correlated to the dysbiosis caused by *Nosema* infection.

**Fig 1.**
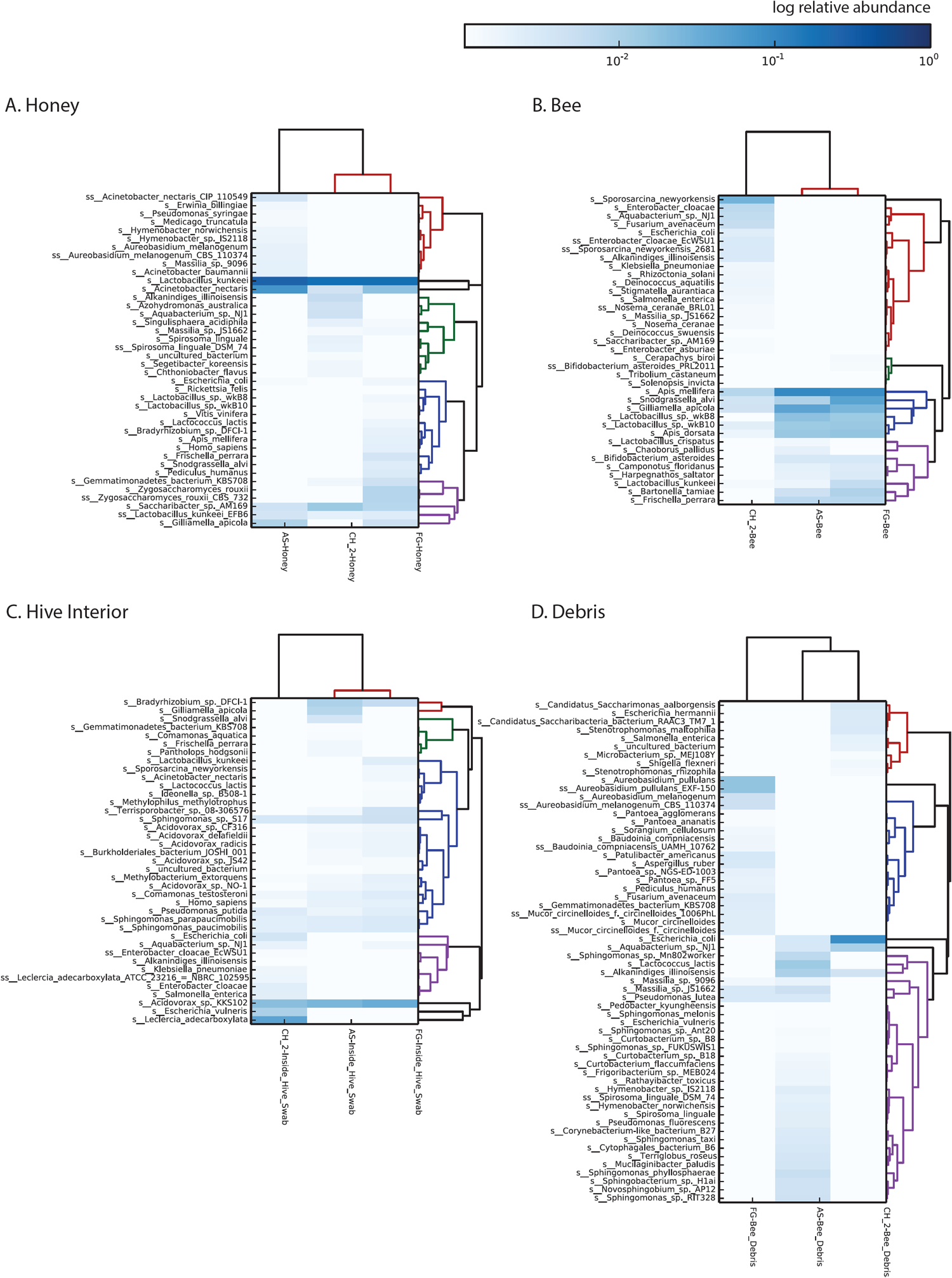
Species classification by type of sample in Brooklyn pilot study: A) Honey, B) Bee, C) Hive interior, D) Debris. Hives are abbreviated as: AS (Astoria), CH (Crown Heights), FG (Fort Greene). *Color map scale corresponds to the log of relative abundance in each sample.*

The inside of the hives was quite uniform across locations, and dominated by environmental bacterial species usually described as found in polluted environments. These include *Acidovorax sp. KKS102*, known to degrade biphenyl/polychlorinated biphenyls (PCBs) (Ohtsubo, Maruyama, Mitsui, Nagata, & Tsuda, 2012), *Sphingomonas sp. S17*, found in high-altitude Andean lakes and tolerant to high pH and desiccation. The interior of beehives is coated with propolis, a resinous substance including polyphenols from essential oils and with a pH of 8.5 (Marcucci, 1994). It is a strong antimicrobial, antifungal and antiviral agent (Kujumgieva et al., 1999) and therefore we hypothesize the presence of extremophile bacteria, and their similar distribution across hives, is a result of selection by the chemical properties of propolis. The species identified in the debris samples were the most diverse (**Table 1**), and include several species of plants as well as plant-associated microbes such at the fungus *Aureobasium pullulans*, also an opportunistic human pathogen (Bolignano & Criseo, 2003), aquatic microbes such as the alkane-degrading *Aquabacterium sp. NJ1* (Masuda, Shiwa, Yoshikawa, & Zylstra, 2014) and mammalian microbes such as the opportunistic pathogen *Stenotrophomonas maltophilia* (Brooke, 2012).

**Table 1.** Beta-diversity according to sample type (Bray-Curtis dissimilarity). P-value calculated against 100 random subsamples of a debris sample. Hives are abbreviated as: AS (Astoria), CH (Crown Heights), FG (Fort Greene).

|  | Honey |  |  | mean | P-value |
| --- | --- | --- | --- | --- | --- |
|  | AS | CH | FG |  |  |
| AS | 0.00 | 0.81 | 0.63 | 0.71 | 0 |
| CH | 0.81 | 0.00 | 0.70 |  |  |
| FG | 0.63 | 0.70 | 0.00 |  |  |

|  | Bee |  |  | mean | P-value |
| --- | --- | --- | --- | --- | --- |
|  | AS | CH | FG |  |  |
| AS | 0.00 | 0.87 | 0.21 | 0.65 | 0 |
| CH | 0.87 | 0.00 | 0.87 |  |  |
| FG | 0.21 | 0.87 | 0.00 |  |  |

|  | Hive Interior |  |  | mean | P-value |
| --- | --- | --- | --- | --- | --- |
|  | AS | CH | FG |  |  |
| AS | 0.00 | 0.59 | 0.28 | 0.47 | 0 |
| CH | 0.59 | 0.00 | 0.54 |  |  |
| FG | 0.28 | 0.54 | 0.00 |  |  |

|  | Bee Debris |  |  | mean | P-value |
| --- | --- | --- | --- | --- | --- |
|  | AS | CH | FG |  |  |
| AS | 0.00 | 0.76 | 0.81 | 0.82 | 0 |
| CH | 0.76 | 0.00 | 0.89 |  |  |
| FG | 0.81 | 0.89 | 0.00 |  |  |

While samples from different hives within a sample type are significantly different from each other (P=0.0) according to Bray-Curtis dissimilarity (**Table 1**), we found the debris samples to be the most diverse, as well as have the highest proportion of environmental bacteria. As our interest was to collect metagenomic information of the environment the bees traverse, rather than that of their hive, we concluded that bee debris is the best material for that purpose.

### 2.2 Urban metagenomes as seen by bees

We next sampled bee hive debris from four cities across the world: Venice, Italy; Sydney and Melbourne in Australia; several neighborhoods in Tokyo. Over all of these locations, we recovered DNA from plants, mammals, insects, arachnids, bacteria and fungi. Taken together, 53% of the classified reads were from multicellular organisms, and 47% from microorganisms. (**Fig 2**).

**Fig 2.**
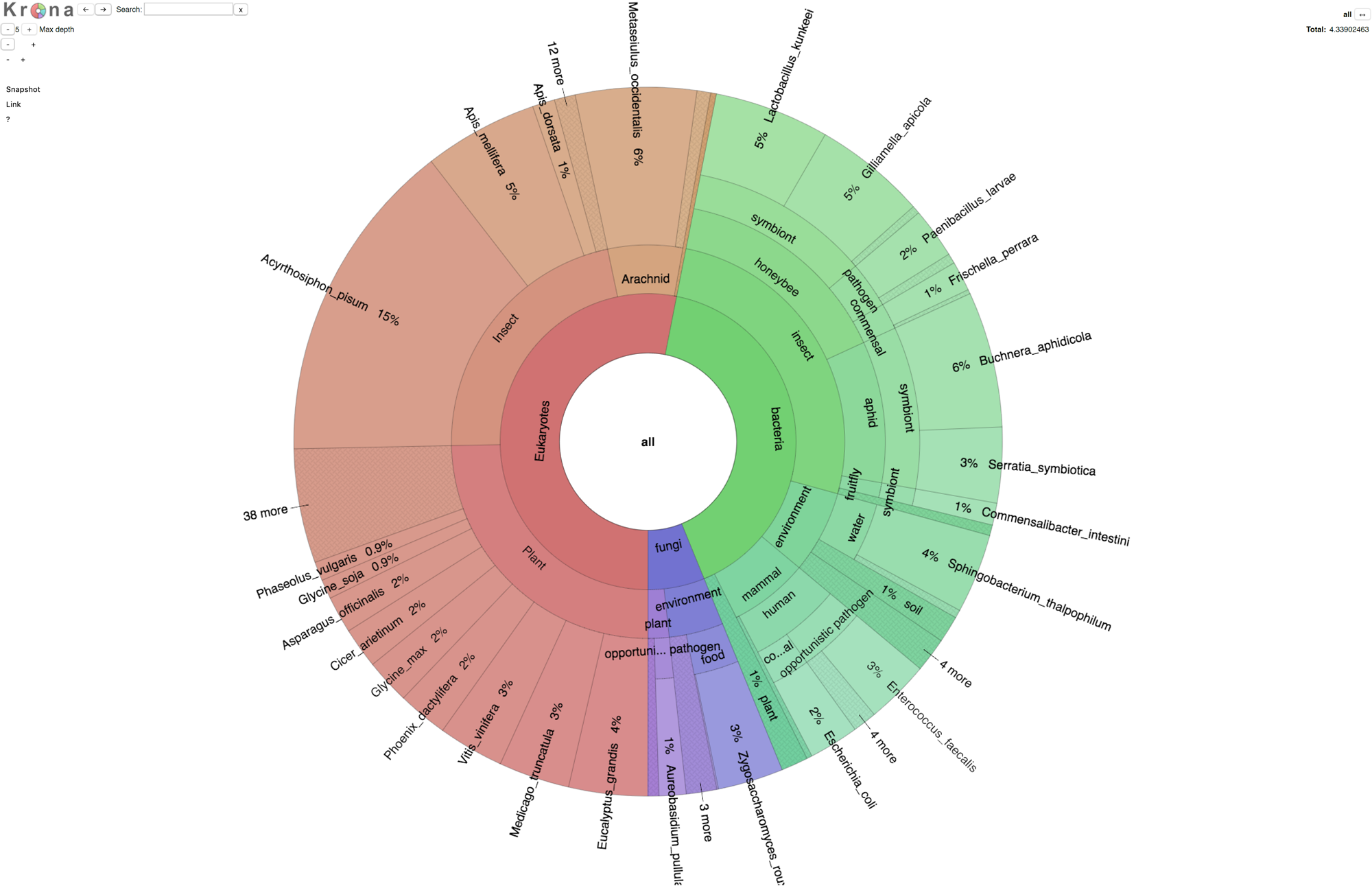
Distribution among kingdoms of classified reads across all samples, with most abundant species in each category.

All metagenomes are significantly different from each other (**Table S1**), and have particularities that can be related to the identity of the city. The metagenome of the Venice debris collected from that hive was largely dominated by fungi related to wood rot (**Fig S2**), which is a common feature of the buildings, built on submerged wooden pilings, and also date palm DNA. Melbourne’s sample was dominated by Eucalyptus DNA, while Sydney’s showed little plant DNA, but bacteria such as *Gordonia polyisoprenivorans*, which degrades rubber (Linos, Steinbuchel, Sproer, & Kroppenstedt, 2017). Tokyo’s metagenome includes plant DNA from Lotus and wild soybean, as well as the soy sauce fermenting yeast *Zygosaccharomyces rouxii*. Overall, each city has a unique metagenomic signature as viewed by bees, with microbes coming from a variety of sources: environmental, insect-related, mammalian and aquatic (**Table 2**).

**Table 2.**
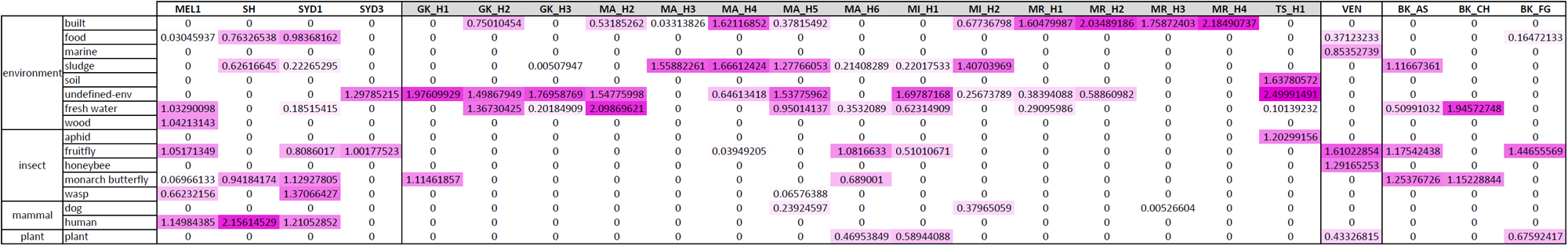
Major classes of bacteria across samples.

### 2.3 Debris as indicator of hive health

As the debris include parts of bees, we looked to the data to see if we could find microbes related to bee health. We found bee commensals (**Table 3**) (Engel, Martinson, & Moran, 2012) as well as potential bee pathogens, namely *Paenibacillus larvae* and the parasite *Varroa destructor*. These results indicate that debris may be used to assess overall hive health.

**Table 3.**
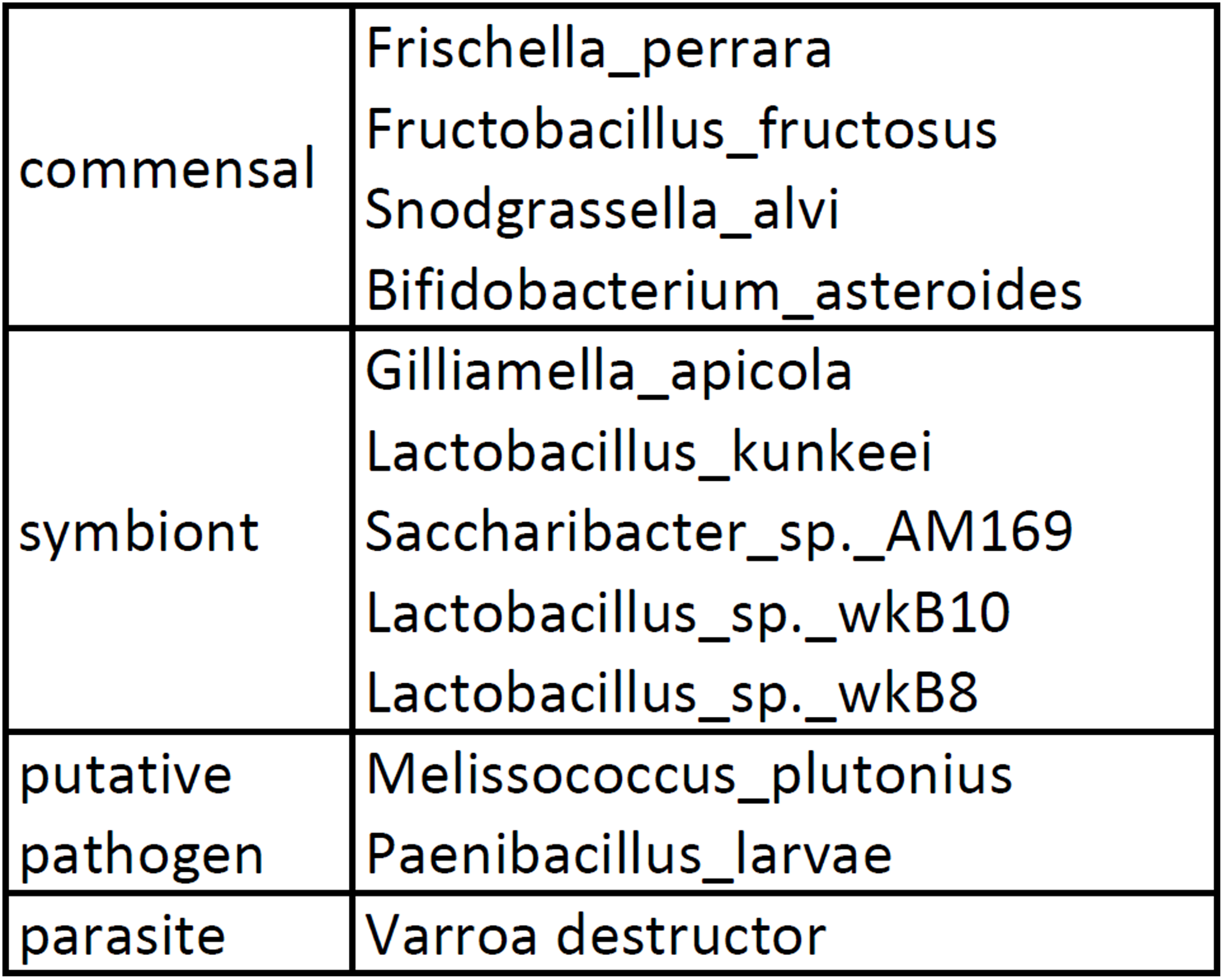
Bee health indicators: known commensals, symbionts, putative pathogens and parasites.

### 2.4 Debris as indicator of human health

As the bees are traversing densely populated urban areas, we tested the hypothesis that they may be able to recover human pathogens. We found various opportunistic pathogens as well as some known disease-causing pathogens, including several species in the Rickettsia genus (**Table 4**). We did de-novo co-assembly of the sequences from a given city to assess the genome completeness of the Rickettsia species as well as the presence of virulence factors. In the Tokyo dataset, we recovered 28 of the 31 *Rickettsia felis* virulence genes at high similarity on the nucleotide level (**Table 5**).

**Table 4.**
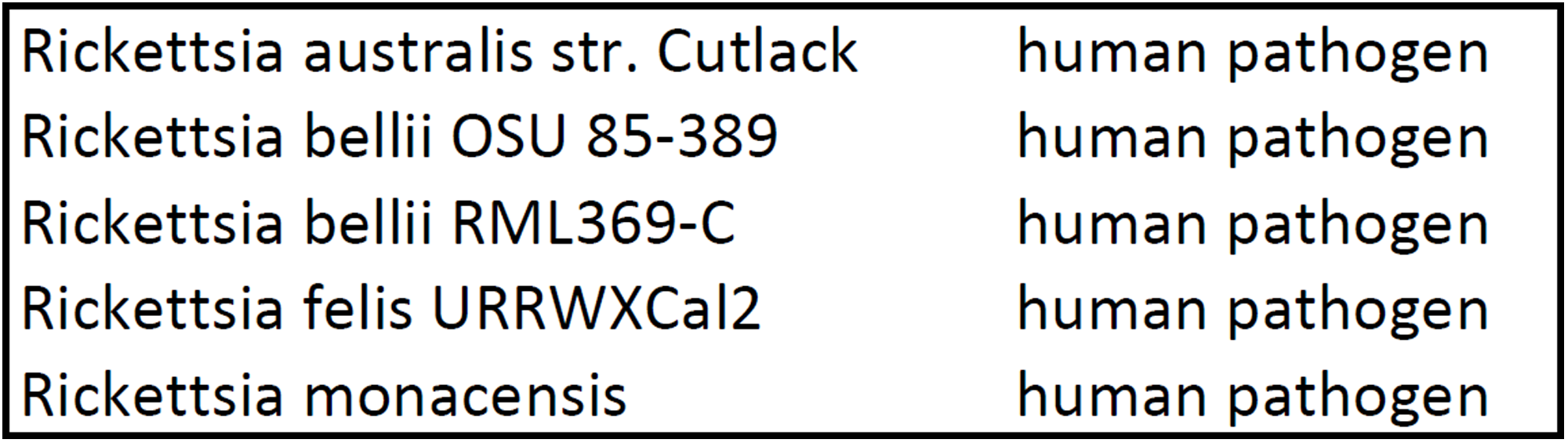
Species of Rickettsia in Tokyo metagenome

**Table 5.** Rickettsia felis virulence factors in assembled contigs of Tokyo metagenome

| gene name | gene length | Alignment %<br>identity | Alignment %<br>gene coverage | Subject contig | Subject contig<br>length |
| --- | --- | --- | --- | --- | --- |
| Accession\|YP_247329adr1 | 870 | 93.02 | 82.07 | k99_3461957 | 17030 |
| Accession\|YP_247330adr2 | 684 | 98.68 | 100.00 |  |  |
| Accession\|YP_246376pat1 | 1650 | 98.37 | 89.33 | k99_5184779 | 4409 |
| Accession\|YP_246759pat2 | 1113 | 98.75 | 101.08 | k99_6299312 | 21131 |
| Accession\|YP_247313pld | 603 | 97.5 | 99.50 | k99_2470097 | 13853 |
| Accession\|YP_246601ralF | 1950 | 97.45 | 100.67 | k99_802869 | 12965 |
| Accession\|YP_246387rickA | 1590 | 97.92 | 66.42 | k99_4497423 | 11956 |
| Accession\|YP_246483rvhB10 | 1446 | 98.8 | 98.27 | k99_2250851 | 6383 |
| Accession\|YP_246484rvhB11 | 1005 | 98.31 | 100.00 |  |  |
| Accession\|YP_246819rvhB1 | 840 | 98.1 | 100.00 | k99_1949946 | 19589 |
| Accession\|YP_247091rvhB2 | 360 | 99.72 | 100.00 | k99_588303 | 7843 |
| Accession\|YP_246103rvhB3 | 288 | 96.88 | 100.00 | k99_2306863 | 24667 |
| Accession\|YP_246104rvhB4a | 2418 | 99.21 | 100.00 | k99_9338833 | 19842 |
| Accession\|YP_247266rvhB4b | 2433 | 99.14 | 100.00 | k99_758254 | 54842 |
| Accession\|YP_246105rvhB6a | 3417 | 98.1 | 100.00 | k99_9338833 | 19842 |
| Accession\|YP_246106rvhB6b | 2019 | 98.09 | 95.99 |  |  |
| Accession\|YP_246107rvhB6c | 2934 | 98.29 | 99.73 |  |  |
| Accession\|YP_246108rvhB6d | 2673 | 98.13 | 100.22 |  |  |
| Accession\|YP_246109rvhB6e | 3468 | 98.7 | 100.00 |  |  |
| Accession\|YP_246480rvhB7 | 180 | 99.44 | 100.00 | k99_7282718 | 13219 |
| Accession\|YP_246479rvhB8a | 699 | 98.86 | 100.00 |  |  |
| Accession\|YP_246481rvhB8b | 732 | 99.18 | 100.00 |  |  |
| Accession\|YP_246478rvhB9a | 753 | 98.94 | 100.00 |  |  |
| Accession\|YP_246482rvhB9b | 477 | 99.58 | 100.00 |  |  |
| Accession\|YP_246485rvhD4 | 1776 | 98.48 | 100.00 | k99_2250851 | 6383 |
| Accession\|YP_246741sca4 | 3120 | 94.57 | 96.22 | k99_796908 | 24425 |
| Accession\|YP_247201tlyC | 900 | 99.11 | 100.00 | k99_7586727 | 10516 |

We assessed the persistence of virulence markers in the debris by taking samples at a 1-week interval in the Tokyo hives. After the first sampling, the bottom trays were cleaned and debris was collected after a week. In some cases, no markers were observed in the second samples, indicating that the cleaning was effective. In the Marunouchi hive H2, markers were found again, and more abundantly (**Table 6**). This indicates virulence markers that are either very abundant in the bee’s range or that they can change rapidly in abundance.

**Table 6.**
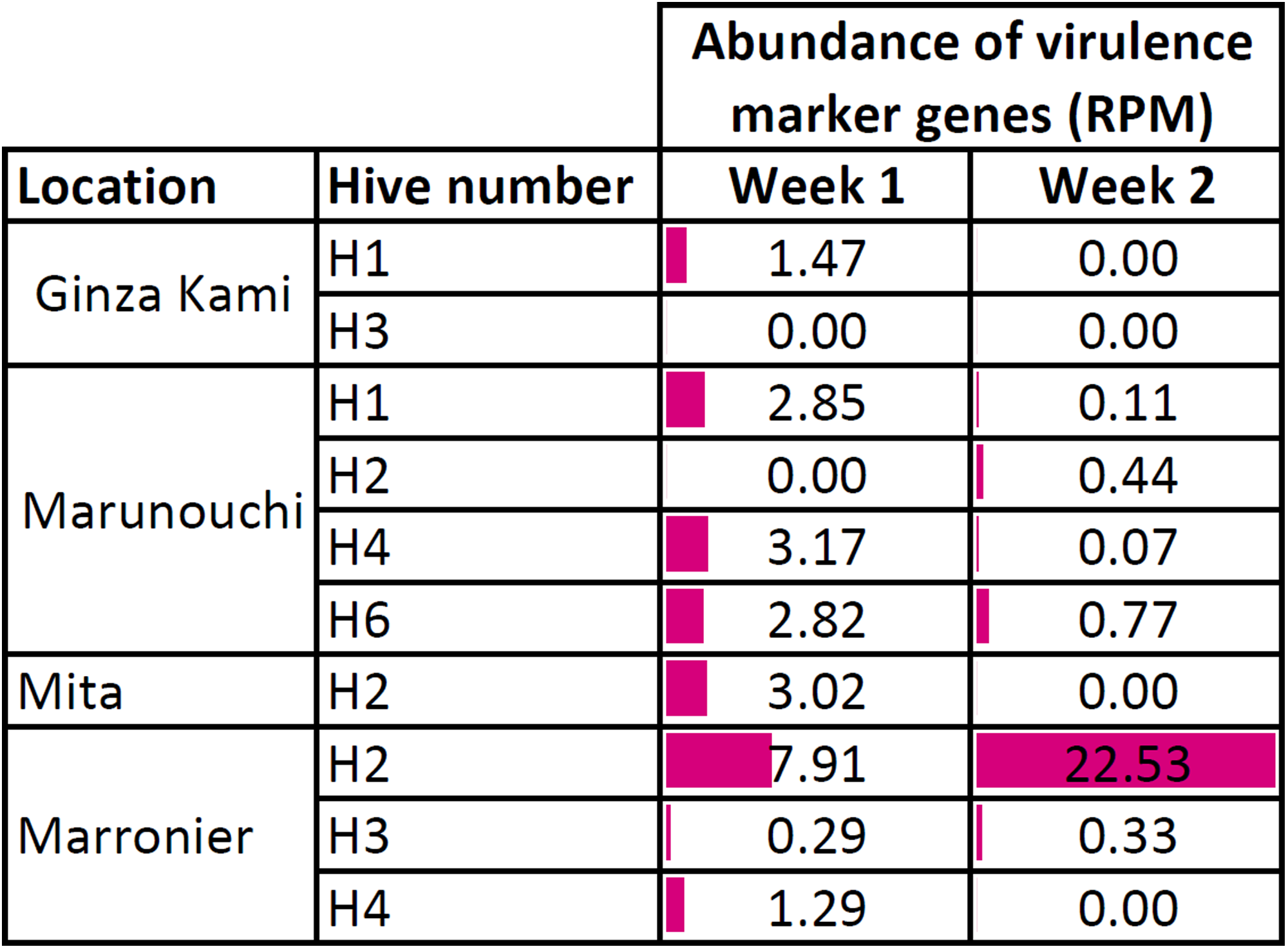
Abundance of virulence factors in at 1-week interval in Tokyo.

|  |  | Abundance of virulence marker genes (RPM) |  |
| --- | --- | --- | --- |
| Location | Hive number | Week 1 | Week 2 |
| Ginza Kami | H1 | 1.47 | 0.00 |
|  | H3 | 0.00 | 0.00 |
| Marunouchi | H1 | 2.85 | 0.11 |
|  | H2 | 0.00 | 0.44 |
|  | H4 | 3.17 | 0.07 |
|  | H6 | 2.82 | 0.77 |
| Mita | H2 | 3.02 | 0.00 |
| Marronier | H2 | 7.91 | 22.53 |
|  | H3 | 0.29 | 0.33 |
|  | H4 | 1.29 | 0.00 |

## 3. DISCUSSION

Here we show that honeybees are relevant sensors for the urban microbiome, and that the debris collected contain a trace of the microbial clouds the bees are traversing. Indeed, these bees recover microbes associated with plants, with which they have physical interactions, but also of mammals and aquatic environments, with which they presumably do not have direct contact. This implies that these microbes were constituents of the respective “microbial clouds” (Meadow et al., 2015) of these entities and that the bees collect a trace of these clouds. Biological content in the atmosphere - the biosphere - was first described in 1978 (Imshenetsky, Lysenko, and Kazakov 1978) and has since been characterized as an integral part of ecosystem function (see Burrows et al. 2009 for review). The biosphere is an indicator of climate change, for example, increasing frequency of dust storms from the African continent are carrying plant and aquatic pathogens to the Americas, affecting coral populations (Behzad, Mineta, and Gojobori 2018). Urban aerosols contain a diverse microbial component including species of potential health and bioterrorism concern. This study demonstrates a novel sampling methodology, which reveals that different neighborhoods have different clouds just as different humans do, and that the collected microbiome can reveal information about the built environment and its inhabitants. For example, the Venetian bees carried a signature of wood rot and aquatic species, similar to previous work showing how flooded areas of a city can carry a “molecular echo” of the aquatic events of its past (Afshinnekoo et al., 2015). Indeed, it has been shown that microbial communities can serve as quantitative geochemical indicators (Smith et al., 2015) and the metabolic properties of the recovered communities can yield information about the environment. Furthermore, metagenomic data can be mined for human-health related information (Rosenfeld et al., 2016). Our ability to recover known bee pathogens as well as virulence factors associated with typhus indicates that this method can serve for early detection of bee-related diseases, and possible human-associated pathogens, in an unprecedented manner. Indeed, insect-based, city-wide microbial monitoring is likely more spatially comprehensive, even if lower resolution, when compared to discrete, human-based sampling techniques, such as swabbing or air-sampling. While the molecular and computational methods used here were based on DNA analysis, it is possible they could be used to monitor RNA-based viruses such as Sars-Cov-2.

In *The Death and Life of Great American Cities* Jane Jacobs describes the city as an ecology:

> “A natural ecosystem is defined as “composed of physical-chemical-biological processes active within a space-time unit of any magnitude.” A city ecosystem is composed of physical-economic-ethical processes active at a given time within a city and its close dependencies”

A city is thus the result of the interactions of its inhabitants, across all species. Our work presented here, and the field of urban metagenomics at large, suggests that the built environment is both a natural ecosystem as well as a city ecosystem in the way Jacobs defines it. We refine this notion then to include not only the interaction of its human inhabitants, but their interactions with their symbiotic microbes. These, taken together, define the identity of a city.

Lynn Margulis in *Symbiosis as a Source of Evolutionary Innovation* (1991) coined the term holobiont, which reframes the identity of eukaryotic, multicellular organisms -- plants and animals -- as more than isolated individuals. Instead, holobionts are “biomolecular networks composed of the host plus its associated microbes” and she suggests that evolution can only be understood under this interaction-based model. In her 2003 pamphlet, “*The Companion Species Manifesto*,” Donna Haraway explores this concept in the context of human-canine relationships and concludes that by inviting these canine species into our lives, humans have inadvertently affected our own human evolution. Thus, the decisions to modify our environment can have influenced the evolution of our species. Although the “*Companion Species Manifesto*” never explicitly points to microbial evolution as a “companion species”, her reasoning can be applied to the metagenome of the city: our design decisions are sculpting a living component of our environment, which in turn influences our health and well-being and ultimately, evolution as a species. The role of city planners and architects, then, would be to design for this living metric.

We have the unique possibility to alter our built environment and design it, not just for ourselves but for all its inhabitants, from environments as common and open as subways (Danko et al, 2019) to those as unique and closed as space stations (Singh et al., 2018, Garrett-Bakelman et al., 2019). As Jane Jacobs says, “Cities are an immense laboratory of trial and error, failure and success, in city planning and city design.” Through studies such as the one presented here, we aim to further understand this accidentally engineered experiment of our built, shared, environment. Some results of this work were shown in physical installation form at the 2016 Venice Architecture Biennale and we believe that the nature of these investigations merit that they be presented in avenues both cultural and scientific.

## 4. METHODS

### 4.1 Hives and collection methods

#### U.S.A. - Brooklyn

The hives of three independent beekeepers were sampled in New York City. The first location (BK1) were Langstroth hives located in Astoria, Queens, NY. The second location (BK2) were Langstroth and Top Bar hives located in Crown Heights, Brooklyn, NY. The third location (BK3) were Langstroth hives located in Fort Greene, Brooklyn, NY. Samples of honey, bees, bee debris, and swabs of the inside of the hive were collected using sterile one-time-use scrapers and transferred into sterile 50ml Falcon tubes.

#### Australia

Two Langstroth hives in Sydney (SYD1, SYD2) and two in Melbourne (MEL, SH) were sampled. Custom collection trays with self-sealing apertures were developed and fabricated at MIT, and shipped to Sydney and Melbourne for deployment. Trays were installed for 1 week collections, then removed and samples were transferred to sterile 50ml Falcon tubes.

#### Italy – Venice

One Langstroth hive at the Palazzo Mora, Venezia, Italy was sampled. Debris were collected from the hive using a sterile one-time-use scraper and transferred to 50ml Falcon tube.

#### Japan – Tokyo

Samples were collected from 12 hives distributed over 4 neighborhoods. Samples were collected with sterile one-time-use scrapers and stored in sterile 50ml

#### Falcon tubes

The locations were Marunouchi (MA), 丸の内 千代田區東京 100-0005, Mita (MI), 港區東京 108-0073日本, Marronnier Gate (MR), マロニエゲート銀座1, and Ginza (GK), 銀座 中央區東京 104-0061.

### 4.2 Sample Preparation

The general approach to DNA extraction involved a combination of lysis methods including mechanical, thermal, and enzymatic disruption to try and ensure that DNA from plant, microbe, and human sources would be extracted for sequencing.

#### Honey

The honey samples were diluted in a 1:1 ratio of grams of honey to mL of ultrapure water and then vortexed vigorously. The mixture was then spun down in the centrifuge at 3900 RPM for 20 minutes, the supernatant was discarded and the pellet+ ∼200 uL residual liquid was moved to an Eppendorf, and placed in the −20°C freezer until the DNA extraction step.

#### Bee Debris

The bee debris was diluted in a 1:5 ratio of grams of bee debris to mL of ultrapure water. The mixture was then heated in a water bath at 70°^C^ for 5 minutes in order to soften the debris and have it disperse in the liquid and then spun on the vortex vigorously. The liquid and solids were then separated, and both were placed into Eppendorfs and placed in the −20°C freezer so that a freeze-thaw cycle would help disrupt the cell membranes. The bee debris material was then ground with a mortar and pestle to break down any large pieces of bee debris, and resuspended in 1X PBS to bring all of the tubes to a final volume of 20 mL. Then material was then allowed to settle, spun down at 39000 RPM for 20 minutes along with 1-2 grams of 100∝m glass beads to further mechanically disrupt the samples. The pellet and a small amount of the supernatant was then used for DNA extraction.

#### Bees

The isopropyl alcohol was drained from the tubes, then bees were placed in a mortar and pestle that was pre-chilled to −80°C before use. The bees were crushed vigorously into a paste. The paste was then placed in Eppendorf tubes and placed in the - 20°^C^ freezer until the DNA extraction step.

#### Swabs

The swabs, Copan Liquid Amies Elution Swab 481C, were stored in the −20°C freezer until the DNA extraction step.

### 4.3 DNA extraction

The protocol for 3-5 mL of starting material of the Promega Wizard® Genomic DNA Purification Kit (A1120) was used, with the following alterations to the standard protocol: one hour incubation at 37°C in a shaker after the neutralization step; the samples were vortexed vigorously for about 1-2 minutes after the lysis and neutralization buffer were added to mechanically disturb the material; following this a phenol/chloroform step was done to remove any remaining organic matter before being placed in the spin column; the DNA was eluted with 20 uL of TE buffer warmed to 65°C; there was a 2 minute incubation time at room temperature before spinning down.

### 4.4 DNA Quantification

The samples were quantified with a Qubit Fluorometric Quantitation device at the Weill Cornell labs, using 1ul input of purified DNA and following manufacturer’s instructions.

### 4.5 Library Preparation

The library preparation followed was developed by Jorge Gandara at the Mason Lab Weill Cornell. It was used to library prep and barcode all samples.

Illumina/Qiagen 500bp Prep:

1. Size selection with Agencourt AMPure XP Beads (A63881)
2. End repair and A-tailing: Qiagen GeneRead DNA Library I Core Kit (180432)
3. Amplification: Qiagen GeneRead DNA Library I Amp Kit (180455)
3. Illumina TruSeq DNA LT adapter kits A and B for up to 24-plex per sequencing pool.

### 4.6 Sequencing

Brooklyn Pilot Study: The samples were sequenced at the BioMicro Center at MIT. The sequencing requested was a 150bp paired end sequence on one lane of the Illumina MiSeq. Venice Study: The sample was sequenced at the CNAG supercomputing center in Barcelona, Spain, with 150bp paired end reads on a Illumina MiSeq lane. Australia and Tokyo samples: Sequencing was performed on the Illumina HiSeq platform at Weill Cornell Medicine, with 125bp paired-end reads. See Supplementary Table 1 for read counts for all samples.

### 4.7 Analysis

#### Metagenomic classification

Read quality was assessed with FastQC and read quality was sufficient to not require trimming. DIAMOND-MEGAN against the NCBI-nr database was used for read classification, as described in McIntyre et al 2017 and McIntyre et al 2019.

run diamond:

~~~
for file in *.fastq.gz; do name=${file/.fastq.gz/}; diamond blastx
-d /path/to/NCBI_nr/nr -q $file -a $name -p 16
~~~

convert binary DIAMOND format to BLAST tabular format:

~~~
for file in *.daa; do diamond view --daa $file --out
${file/.daa/}.tab --outfmt tab; echo $file; done
~~~

perform read-by-read taxonomy classification with MEGAN:

~~~
for file in *.tab; do /path/to/programs/megan/tools/blast2lca --
input $file --format BlastTAB --topPercent 10 --gi2taxa
/path/to/programs/megan/GI_Tax_mapping/gi_taxid-March2015X.bin --
output $file.read_assignments.txt; done
~~~

Heatmap in Figure 1 was generated with the script `metaphlan_hclust_heatmap.py` from the MetaPhlan package, displaying the abundances for species only (default `-- tax_lev s`), in logarithmic scale `(-s log`). The clustering is performed with “average” linkage (default `-m average`), using “Bray-Curtis” distance for clades (default `-d braycurtis`) and “correlation” for samples (default `-f correlation`).

~~~
metaphlan_hclust_heatmap.py --in $file --out
$file.Blues.minv0.maxv1.Blues.log.pdf -c Blues -s log --minv 0.0 --
maxv 1
~~~

#### Diversity quantification

Beta-diversity was calculated according to the Bray-Curtis dissimilarity metric (Bray and Curtis 1957) as implemented by the Qiime package (Caporaso 2010, http://qiime.org/).

~~~
$ metaphlan2biom.py merged.samples.metaphlan.out
merged.samples.biom
$ beta_diversity.py -i merged.samples.biom -m bray_curtis -o
merged.samples.beta_div.bray_curtis
~~~

P-value was calculated based on 100 bootstrapped subsamples of the Brooklyn debris sample, each subsample being of 1 million reads. Bootstrapped samples were classified using same methods as described above, and pairwise beta-diversity calculated as above. P-value was calculated as the number of bootstrap samples with lesser dissimilarity value than the test value.

#### Assembly

Co-assembly of Tokyo samples (assembly of all sequences pooled together) was performed with MegaHit (Li, Liu, Luo, Sadakane, & Lam, 2015) and reads for each individual sample were mapped to contigs with Bowtie2 (Langmead & Salzberg, 2012). Assembly yielded 3207501 contigs, see Supplemental Table 2 for assembly statistics.

## Supporting information

SRA metadata

## 5. ACKNOWLEDGEMENTS

DN and MP located the hives and took samples. MP designed custom trays for sampling. DN developed specific DNA extraction protocols for bee material. EH and DN prepared libraries for sequencing and EH analyzed the data. RF and MP developed the data visualization and MP and CW designed the installation for the Venice Biennale exhibit. KS and CM supervised the project and helped design experiments and provided logistical support. EH drafted the manuscript, with help from RF and DN and edits from CM. All authors have read and approved the manuscript. The authors would like to acknowledge the Mori Building Company for their financial support, and Jun Fujiwara especially for his continued interest in the project. We would like to thank all of the beekeepers for so generously supporting this project with their time and materials. We would also like to thank Mike Laserwalker and Ben Berman for their assistance in the Biennale exhibit, as well as Daniela Bezdan for her support in sequencing preparation. CEM would like to thank the Epigenomics Core Facility, the Vallee Foundation, Igor Tulchinsky and the WorldQuant Foundation, the National Institutes of Health (1R01MH117406), the Bill and Melinda Gates Foundation (OPP1151054), the NSF (1840275), and the Alfred P. Sloan Foundation (G-2015-13964).

---

**Supplementary Table 1.**

|  | Sample | Million Reads |
| --- | --- | --- |
| Brooklyn | AS-Bee | 1.519066 |
|  | AS-Bee_Debris | 1.690898 |
|  | AS-Bee_Debris | 1.918464 |
|  | AS-Honey | 1.754984 |
|  | AS-Inside_Hive_Swab | 1.14609 |
|  | CH_2-Bee | 1.823688 |
|  | CH_2-Bee_Debris | 1.865624 |
|  | CH_2-Bee_Debris | 1.368362 |
|  | CH_2-Honey | 1.72086 |
|  | CH_2-Inside_Hive_Swab | 1.282562 |
|  | FG-Bee | 1.819204 |
|  | FG-Bee_Debris | 1.907648 |
|  | FG-Honey | 1.255524 |
|  | FG-Inside_Hive_Swab | 1.487172 |
| Tokyo | GK_H1_S1_S12 | 12.22419 |
|  | GK_H1_S2_S9 | 11.449704 |
|  | GK_H3_S1_S10 | 10.703352 |
|  | GK_H3_S2_S8 | 28.83604 |
|  | MA_H1_S1_S22 | 58.925544 |
|  | MA_H1_S2_S2 | 130.585438 |
|  | MA_H2_S1_S21 | 21.018622 |
|  | MA_H2_S2_S1 | 13.770676 |
|  | MA_H4_S1_S19 | 24.612872 |
|  | MA_H4_S2_S17 | 26.797518 |
|  | MA_H6_S1_S3 | 0.70942 |
|  | MA_H6_S2_S16 | 23.242922 |
|  | MI_H2_S1_S14 | 17.856454 |
|  | MI_H2_S2_S13 | 17.317142 |
|  | MR_H2_S1_S4 | 14.153042 |
|  | MR_H2_S2_S27 | 25.658876 |
|  | MR_H3_S1_S29 | 27.467216 |
|  | MR_H3_S2_S26 | 30.420888 |
|  | MR_H4_S1_S28 | 23.282586 |
|  | MR_H4_S2_S25 | 12.263354 |
| Australia | MEL1 | 46.579378 |
|  | SH | 36.062846 |
|  | SYD1 | 42.08362 |
|  | SYD3 | 45.179888 |
| Venice | Venice | 24.690872 |

**Supplementary Table 2.**

| **number of**<br>**contigs** | **total bp** | **min contig**<br>**length** | **max contig**<br>**length** | **average**<br>**contig length** | **N50** |
| --- | --- | --- | --- | --- | --- |
| 3207501 | 2802811167 | 200 | 488034 | 874 | 1515 |

